# Rapid expansion overcomes inbreeding in cross-hemisphere colonization of Barn Swallows

**DOI:** 10.64898/2026.07.13.738246

**Authors:** Valentina Gómez-Bahamón, Juan I. Areta, Jeremy Summers, Cristian Torres, Facundo Gandoy, Mingzuyu Pan, Zachary A. Szpiech, David P. L. Toews

## Abstract

Switching migratory behavior appears to promote speciation in birds through colonization of their wintering grounds for breeding. However, colonization episodes often fail, especially when they lead to inbreeding depression associated with small population sizes. We studied the dynamics of a new and thriving breeding population of migratory Barn Swallows (*Hirundo rustica erythrogaster*) that began nesting on their traditional wintering grounds in South America in the 1980’s and evolved an “inverse” migration and breeding cycle. We show that in four decades, behavioral flexibility and standing genetic variation have facilitated the establishment of a breeding population that has grown exponentially in abundance and breeding range size. By breeding under bridges and culverts, Barn Swallows exploited ecological opportunities that outweighed the costs of inbreeding and began an independent evolutionary trajectory from their North American relatives.

## Introduction

Organisms expand into new geographic areas by dispersing and establishing new breeding populations. Long distance dispersal events can involve a small number of colonizing individuals that successfully breed. Such a process, coupled with local adaptation and long periods of isolation, can sometimes lead to the origination of a new species (*1*, *2*). However, the population size reduction associated with colonization episodes can lead to subsequent erosion of genetic diversity via genetic drift, weakening the adaptive potential of populations to persist in new environments and increasing failure likelihood of the colonization attempt (*3*). A decrease in population size also increases the probability of inbreeding, which can unmask recessive deleterious variation, causing inbreeding depression and increasing extinction risk (*4*). A critical stage in the colonization process occurs during the initial growth phase before the population reaches equilibrium, in which natural selection acts on the genetic and ecological attributes of the new population to persist (*5*).

During population establishment, the ecological context encountered by colonizers in a new locality affects both the probability of persistence and of divergence. Colonization events can involve populations dispersing over long distances to breed in geographic localities with novel ecological opportunities and challenges. However, successful establishment following these events depends on a combination of intrinsic factors apart from extrinsic environmental pressures. Specifically, the attributes of the genetic pool from which the population is founded (*6*) and the standing genetic variability (*7*), as well as the behavioral and phenotypic flexibility of the individuals involved in the process (*8–10*), in concert with the encountered ecological context they encounter can enable or hamper successful colonization.

Empirically understanding the intrinsic and extrinsic factors influencing growth rate can shed light into which attributes contribute to population persistence after a natural colonization episode in the wild. Yet, following this play out in real-time has been nearly impossible for most systems. The North American Barn Swallow (*Hirundo rustica erythrogaster*), by contrast, provides a unique opportunity to study this process during a natural colonization event. This species uses buildings as support structures to construct their nests, and every year they migrate south to spend the winter in South America (Fig. 1A) (*11*), involving a migratory journey of ∼4,000-11,000 kilometers (*12*). In 1980, a small group of Barn Swallows was observed nesting under a bridge in the Buenos Aires Province, and hatched nestlings from 6 successful nests (Fig. 1B) (*13*). This was the first observation of the species successfully breeding in the southern hemisphere. Since that time, ornithologists in South America have monitored and reported the presence of nests as this population rapidly expanded, today reaching both the Atlantic and Pacific coasts of southern South America, breeding under bridges and culverts (Fig. 1C) (*13–29*), and more recently in buildings as well. The population has also evolved an inverse migratory cycle, migrating to northern South America during the Austral winter (*30*, *31*), associated with a significant reduction in the overall migratory distance. We studied the dynamics of the interplay between ecological opportunity, behavioral flexibility, and population genetic attributes as a colonization event is happening in ecological timescales, at the new breeding (former wintering) grounds in the opposite breeding hemisphere.

**Fig. 1.**
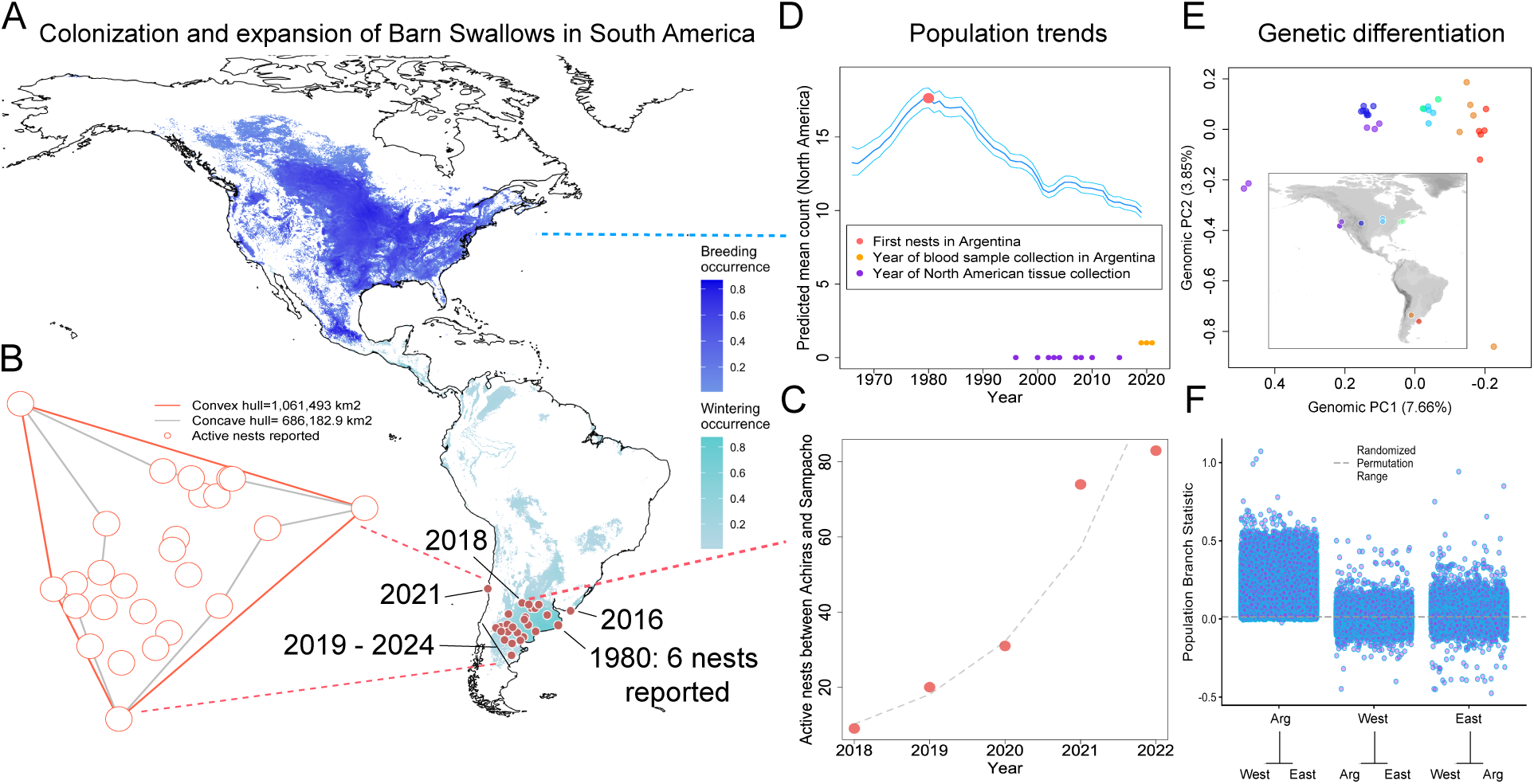
Geographic and demographic evolution of Barn Swallows in South America following a founder event in 1980. **(A)** Breeding and wintering distributions of Barn Swallows in the American continent before and after the first observation of nests in South America in the year 1980. Slate blue range shows the breeding area in North America, turquoise the wintering area of North American breeding Barn Swallows obtained from eBird occurrence data (*51*). Nesting localities in South America are indicated with red circles. **(B)** Estimate of the breeding range size of the recent South American breeding population based on published reports of active Barn Swallow nests. The area within the red lines indicates the convex hull encompassed by those localities and the area in gray indicates the concave hull. **(C)** Five-year change in number of active nests in a transect between Sampacho and Achiras in the Province of Cordoba, Argentina, a starting from the year when the firsts nest was reported there. **(D)** Abundance estimates through time of the North American breeding Barn Swallows obtained from the North American Breeding Bird Survey (*33*). The red circle indicates the time when nests were first reported in Argentina (*13*). The purple circles represent the years in which the museum tissues of the North American breeding birds included in this study were collected. The orange circles indicate the years when we sampled blood from wild caught Barn Swallows in Argentina. **(E)** Genomic Principal component analysis based on genotype likelihoods across the genome. The map indicates the localities where the samples used to generate the PCA were collected. **(F)** Genetic differentiation estimated with Population Branch Statistic across 10 Kb non-overlapping windows of the genome. Argentine birds are the most highly differentiated when compared to North American population together, followed by western North American birds from Argentine and eastern North American, and the least differentiated is Eastern North America, which most closely shares ancestry with both Western North American and Argentine populations.

### Rapid range expansion and increase in abundance of Argentine Barn Swallows

To estimate demographic changes in the new breeding population of Barn Swallows, we first quantified the geographic range expansion based on published reports of nests in different areas of South America (*13–29*) (Figure 1A). We estimated the area encompassed within the outermost nest localities from the convex hull, finding that in 43 years (1980–2023) the population expanded over ∼1,061,500 km^2^ from the original location (Figure 1B). To estimate changes in abundance, we identified an area in Cordoba Province (Argentina), where the Barn Swallows had not begun nesting until the year 2018 (*19*). From that year onwards, we monitored bridges and culverts between the towns of Achiras and Sampacho (70 km away; supplementary Fig. S1) for five consecutive years, recording the number of active nests (i.e., those with incubated eggs or nestlings that were fed by adults). We found that the abundance of active nests increased exponentially by a factor of *r* ∼0.58 (Fig. 1C). To compare these demographic dynamics with those of the North American breeding population, we used a publicly available dataset (The North American Breeding Bird survey), consisting of yearly survey data provided by hundreds of volunteer birders across Canada and the United States since 1966 (*32*). This curated dataset includes estimates of relative abundance (*33*). We found that at the same time as Barn Swallows began breeding in Argentina, the abundance of breeding North American Barn Swallows began to steadily decline (Fig. 1D). Moreover, the population in North America continues to decline, whereas the growth curve in the Cordoba Province in Argentina shows an exponential increase.

### Argentine Barn Swallows are on an independent evolutionary trajectory

To assess whether this founder event has led to genetic differentiation of the Argentine population with respect to North American Barn Swallows, we initially estimated population genetic attributes from whole genome resequencing data (∼6x coverage) for (i) breeding individuals captured at two localities in Argentina (close to the initial colonization area in Buenos Aires and at the western expansion front in Cordoba), and (ii) museum tissues from breeding individuals in Eastern and Western North America (collected between the years 1996 and 2008) (Fig. 1E, supplementary Table S1). We additionally included an independently published dataset of whole genome resequencing data, with similar coverage to ours, from western USA (*34*). Despite their very recent origin, we found that breeding Barn Swallows in Argentina comprise a distinguishable genetic cluster along genomic PC1, with respect to North American breeding birds (Fig. 1E). Moreover, we found that Argentine Barn Swallows cluster closer to the Eastern North American breeding birds than to those from the West, suggesting that the Argentine population was primarily founded by Eastern North American birds. These results are consistent with geolocator tracking data of individual movements showing that eastern North American Barn Swallows migrate to southern South America, whereas the western birds migrate to central and northern South America (*12*). This is also consistent with admixture proportions— assuming 2 genetic clusters (i.e., K=2)—where the Eastern North American breeding birds appear genetically associated with a combination of Western North American and Argentine breeding birds (supplementary Fig. S2A). Furthermore, a distance-based tree supports a monophyletic relationship among the Argentine breeding birds (supplementary Fig. S2B), indicating a single colonization event (with the potential influx of additional migrants) and an independent evolutionary trajectory.

To further quantify genetic differentiation, we estimated population differentiation across the genome with *F*_ST_ and PBS for 10 Kb non-overlapping windows. We found that pairwise comparisons of western and eastern North American populations with the Argentine population exhibited small but significant overall differences in allelic frequencies than the pairwise comparison between western and eastern North American populations (supplementary Fig. S3). Mean *F*_ST_ estimated between eastern and western North America was low (0.04), consistent with high levels of gene flow among them, and what would be expected for birds with the dispersal capabilities of a Barn Swallow (based on their flight morphology (*35*)). By contrast, mean *F*_ST_ between Argentina and western North America was 0.12, and versus eastern North America was 0.09, showing a small but significant increase in differentiation in the Argentine population across the genome after the founder event in 1980. Likewise, population branch statistic shows significantly larger differentiation when Argentina is the focal population (Fig. 1F).

### No evidence of strong positive selection in Argentine Barn Swallows

What intrinsic and extrinsic factors lead to migratory switches are still unknown. One hypothesis is that structural mutations, such as chromosomal inversions, which harbor multiple genes coding for instructions among correlated traits associated with migratory behavior could facilitate transitions to a new migratory program or loss of migration (*36*). To test whether there are regions of the genome associated with the differences in migratory behavior we investigated the variation in allelic frequencies across the genome with *F*_ST_ scans for data with a higher depth coverage (average 11X). Although we found no fixed genetic differences in any comparison between the North American and Argentine Barn Swallows, we found three highly differentiated regions (*F*_ST_ = 0.35-0.47) on chromosome 3 and one on chromosome 6 (*F*_ST_ =0.82) (Fig. 2A). The regions in chromosome 3 are located within the genes *SERTAD2,* associated with fat storage; *LOC131378466,* a myotubularin-like gene of lipid phosphates involved in membrane trafficking and endocytosis; and *LOC120751327,* which has an uncharacterized gene function in birds. The differentiated region on chromosome 6 is on a non-coding region upstream to another uncharacterized gene (*LOC120754407*). Contrary to what would be expected from strong positive selection following a colonization event, none of these regions show divergent values of Tajima’s D between North and South American breeding birds (Fig. 2B and 2C). On the contrary, along both chromosomes, the regions with high *F*_ST_ values appear to be under strong balancing selection (Fig. 2B and supplementary Fig. S4). To deconstruct this further on chromosome 3, we inspected whether *F*_ST_ peaks coincided with peaks in Tajima’s D and with peaks in allelic diversity in both populations (Fig. 2D and 2E). We found that these regions comprised high genetic variation as well as high values of Tajima’s D. These results indicate balancing selection as opposed to selective sweeps driving outlier *F*_ST_ values in the Argentine population.

**Fig. 2.**
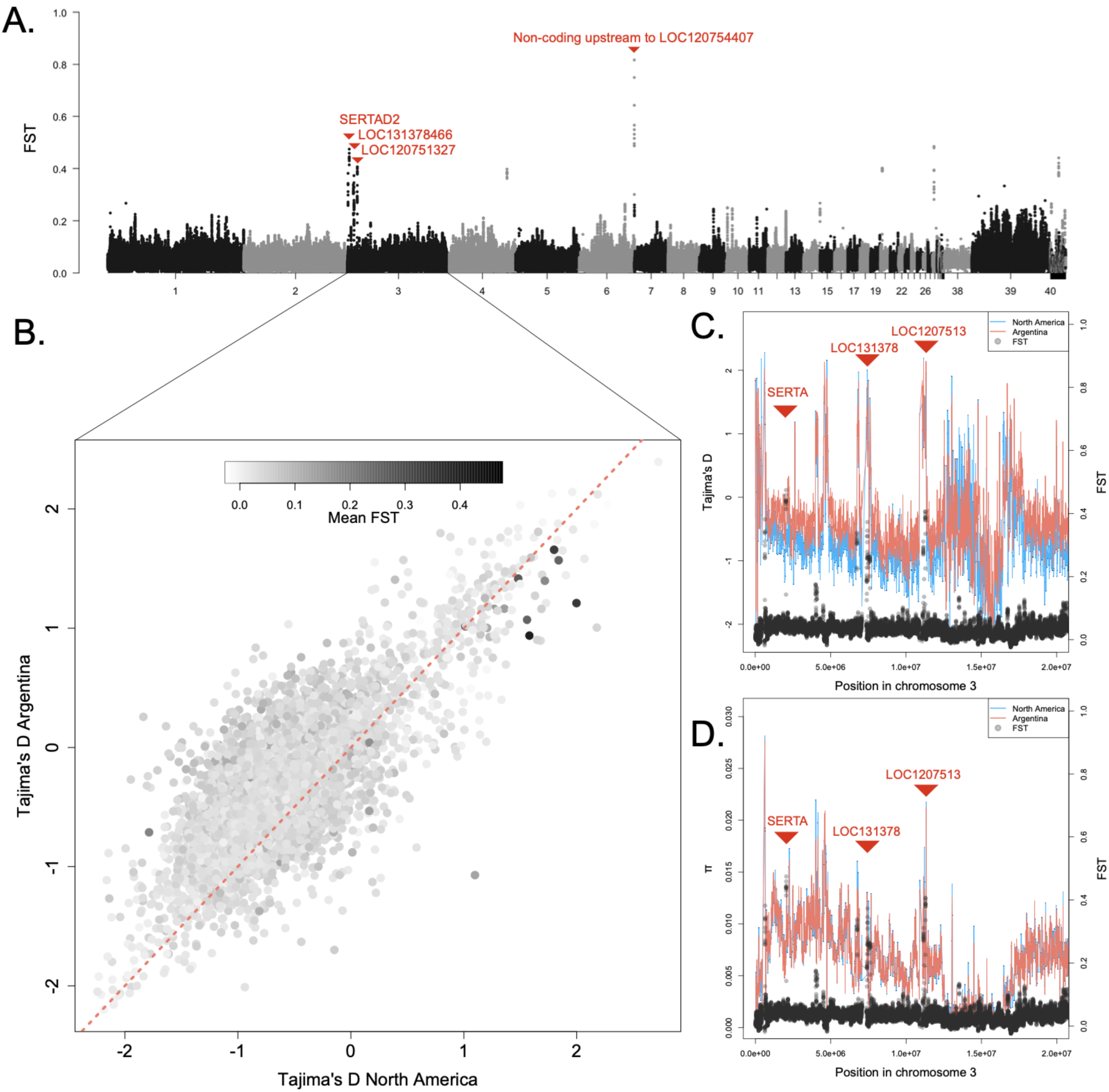
Outliers of genetic differentiation and diversity across the genome of Barn Swallows. **(A)** *F*_ST_ scans between North American and Argentine Barn Swallows showing regions with outlier values of differentiation. Red text indicates whether the outlier regions are located on genes annotated in the Barn Swallow reference genome or in non-coding regions. **(B)** Tajima’s D estimates for North America and Argentina across chromosome 3 where positive values indicate evidence of balancing selection and negative values of selective sweeps. Outlier regions are colored in black corresponding to the estimated *F*_ST_ values. The points closer to the red dotted line indicate evidence for similar processes acting on that region in Argentina and North America. **(C)** Estimates of Tajima’s D for Argentina (salmon) and North America (blue) across the outlier region in chromosome 3 showing that the regions with higher differentiation also show higher values of Tajima’s D in both parental and founder populations. **(D)** Estimates for nucleotide diversity (*π*) cross the outlier region in chromosome 3 showing that the regions with higher differentiation also show higher values of nucleotide diversity.

### The Argentine population of Barn Swallows is strongly inbred

Severe reductions in population size can lead to increased inbreeding. A genomic signature of such inbreeding is that offspring born after the colonization episode tend to have long continuous homozygous regions in their genomes, called runs of homozygosity (ROH). Theoretical expectations under different demographic scenarios are revealed by attributes of these ROH, specifically how many and how long they are in sum (*37*). We estimated ROH attributes in the Barn Swallows from whole genomes of Argentine and North American breeding individuals. We estimated the number and sum of runs of homozygosity, finding that the North American breeding birds match expectations for an admixed population, whereas the South American breeding population has a skewed distribution with individuals having a large proportion of their genome in contiguous homozygosity (Fig. 3A), as expected for a population with inbred individuals (Fig. 3A) (*37*).

**Fig. 3.**
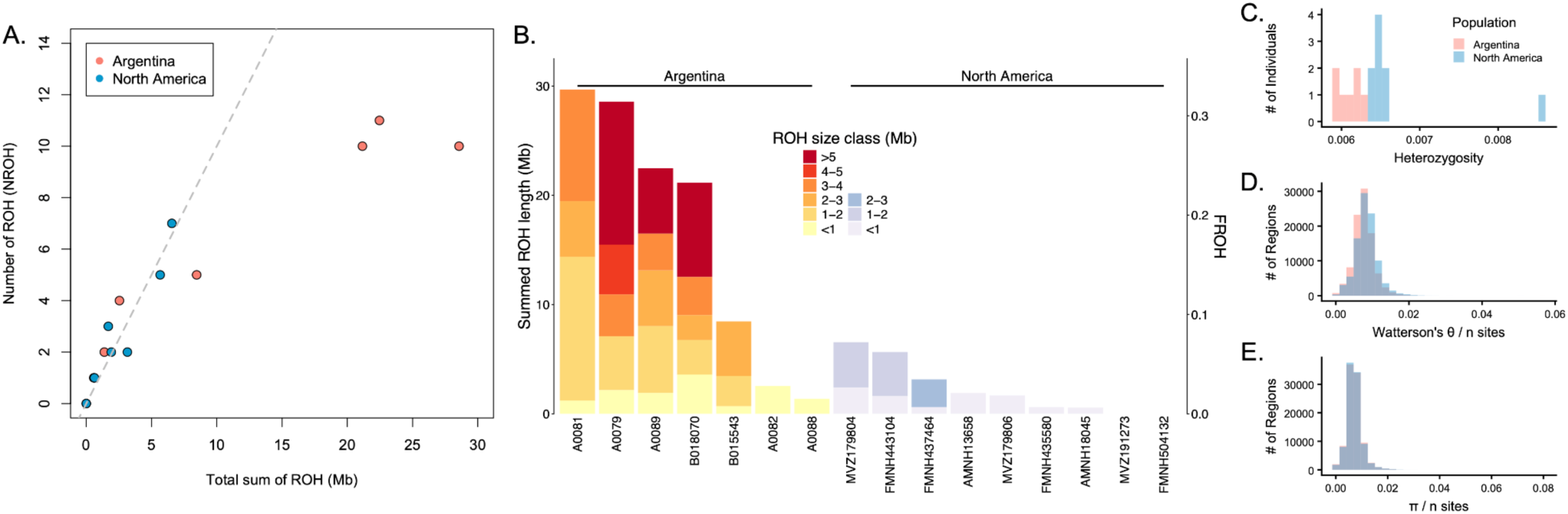
Population genetic attributes of parental and founder populations of Barn Swallows. **(A)** Number and sum of runs of homozygosity (ROH) per individual indicating that there are inbred individuals in the Argentine population (salmon), whereas in North America (blue), all individuals fall within the expectation of an admixed population (gray dotted line). **(B)** ROH by size classes (Mb) found for individuals. FROH indicates the fraction of the whole genome (excluding sex chromosomes) that is under ROH. Labels in the x axis indicate the individual ID. Other measures of genetic diversity estimated for the Argentine (salmon) and North America (blue) populations, including **(C)** heterozygosity, **(D)** Waterson’s *θ* divided by the number of sites, and **(E)** *π* divided by the number of sites.

To further investigate potential evidence of inbreeding in the Argentine population, we compared individuals separated by ROH size classes. We found that the Argentine Barn Swallows have individuals with long runs of homozygosity (>5Mb) comprising >0.3 of their genomes, suggesting some, but not all, individuals are highly and recently inbred, whereas North American birds either lacked ROH or had a maximum of 2-3Mb of continuous sequences in homozygosity (Fig. 3B, supplementary Fig S6). Consistent with this, we also found that the Argentine population exhibited weak but significantly lower individual heterozygosity in comparison to North America (Fig. 3C). We also found a small but significant difference in genetic variability estimated with Waterson’s *θ* with the North American population having slightly and consistently higher values (Fig. 3D). However, nucleotide diversity (*π*;Fig. 3E) showed no significant differences among the populations. One potential explanation is that gene flow could be maintaining allelic diversity. Past work invoked gene flow given little-to-no genetic differentiation based on a small number of microsatellite markers or mitochondrial *ND2* sequences (*38*). Because North American Barn Swallows migrate to South America to overwinter at the same time that the South American Barn Swallows are breeding, there have been recurrent opportunities for gene flow from new North American birds into the genetic pool of South American breeders, after the initial founder event.

### Little demographic connection between North American and Argentine Barn Swallows

To explicitly test whether gene flow has occurred during the establishment of the new South American breeding population, we ran demographic models informed with the population growth curves obtained from the citizen science dataset in North America (*33*), as well as our estimates from surveys in areas of active nesting in Cordoba, Argentina. We tested three scenarios that varied based on whether there was (i) no gene flow during divergence, (ii) bidirectional, or (iii) unidirectional geneflow between North American and Argentine Barn Swallows (Fig. 4A-C). The parameters estimated in these simulations included their initial effective population size, their current effective population sizes, time since divergence, and magnitude of gene flow. We found that all the highest ranked models based on *AIC* comparisons involved bidirectional gene flow, although the estimated values for gene flow under the higher fit models were very small (2E-07 to 5E-07) (Fig. 4C). Assuming a 2-year generation time and a mutation rate of 2.3×10^-9^ mutations per site per year (*39*), the models with higher support estimated a divergence time with a skewed distribution towards divergence time of less than 100 years ago (Fig. 4B). This is consistent with the published natural history observation of establishment in 1980 (*13*). Moreover, higher support for models with gene flow and a recent colonization were also consistent when simulating scenarios that do not constrain population growth rate (Fig. 4C and D).

**Fig. 4.**
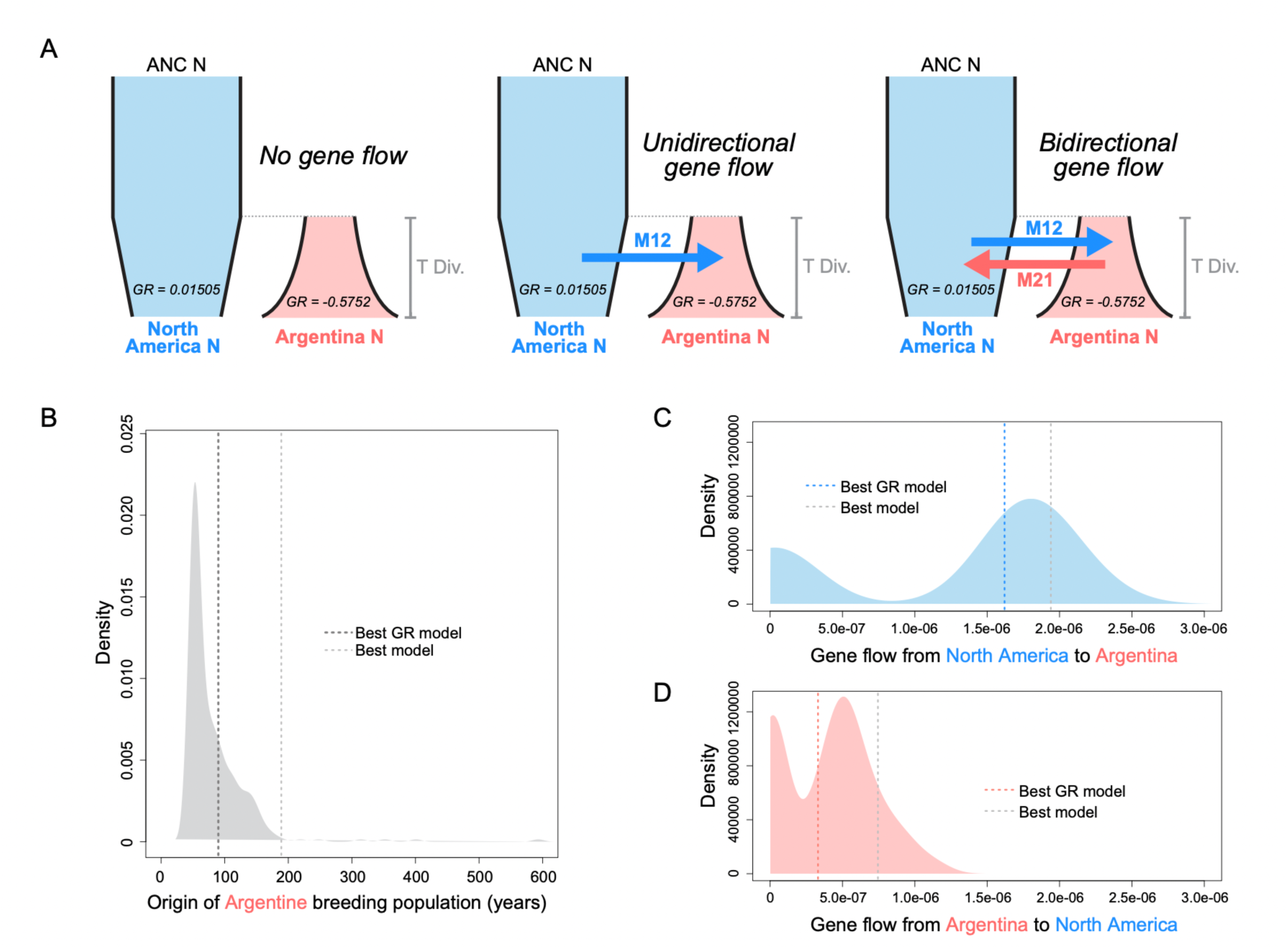
Demographic inference testing model fit of divergence scenarios with and without gene flow in Barn Swallows. **(A)** Models tested using population growth curve estimated from the North American breeding Barn Swallows obtained from the North American Breeding Bird Survey (*33*) and our data collected on active colonies in Cordoba, Argentina. Models differ on whether they allow unidirectional, bidirectional or no gene flow. Models without growth rate were also tested. All highest ranked models with and without constraining growth rates, had bidirectional geneflow. **(B)** The estimated time since divergence for all models peaks around the time when the first nests were observed. Dashed lines indicate the estimated values for the highest ranked models that constrain and do not constrain growth rate. **(C)** The Estimated values for gene flow for all models including models that constrain and do not constrain growth rate from North America to Argentina. Dashed lines indicate the estimated values for the highest ranked models that constrain and do not constrain growth rate. **(D)** The Estimated values for gene flow for all models including models that constrain and do not constrain growth rate from North America to Argentina. Dashed lines indicate the estimated values for the highest ranked models that constrain and do not constrain growth rate.

Speciation in birds has been thought to occur primarily with geographic isolation (*40*), followed by the accumulation of genetic mutations through time that can lead to genetic incompatibilities (*2*). In migratory species that breed in the wintering grounds, ecological speciation has also been shown to promote prezygotic and postzygotic isolation and can lead to rapid speciation (*41*). To understand how the genetic divergence we estimate is brought about in the Argentine Barn Swallows, given the regular presence of migratory individuals and evidence of gene flow, we reconstructed their presumed evolutionary trajectory based on simulated data. Specifically, we aimed to determine the initial population size and amount of gene flow that would be required to reconstruct the observed population genetic attributes estimated with the Barn Swallow data. We first simulated the evolution of a parental population undergoing a demographic decline resembling North American Barn Swallows, with a founder event and exponential growth resembling Argentine Barn Swallows. We then tested nine combinations of founding population sizes (ranging from 6-100) and gene flow (ranging from 0 to 0.01). We estimated genetic diversity and ROH from sampled individuals. We found that the attributes that most closely resembled the empirical data with long runs of homozygosity corresponded to simulations with the smallest initial population size (i.e., assuming 5 female fertilized by one male, totaling 6 individuals) (Fig. 5A). The simulated values of gene flow, on the other hand, did not influence FROH (Fig. 5A). Consistent with our empirical results, heterozygosity and ROH were decoupled. We found that Heterozygosity matched a scenario with the smallest population size— but contrary to FROH—higher values of gene flow increased heterozygosity (Fig. 5B). Nucleotide diversity, on the other hand, matched the larger population size simulated with 100 founding individuals (Fig. 5C). Broadly, our simulated results corroborate the finding that, although a population involves mating among close relatives, other measures of allelic diversity can remain stable when population growth is rapid. The combination of standing genetic variation, gene flow, and small founding population sizes followed by rapid population increase and range expansion can lead to divergence in allelic frequencies, persistence and potentially speciation despite mating between related individuals.

**Fig. 5.**
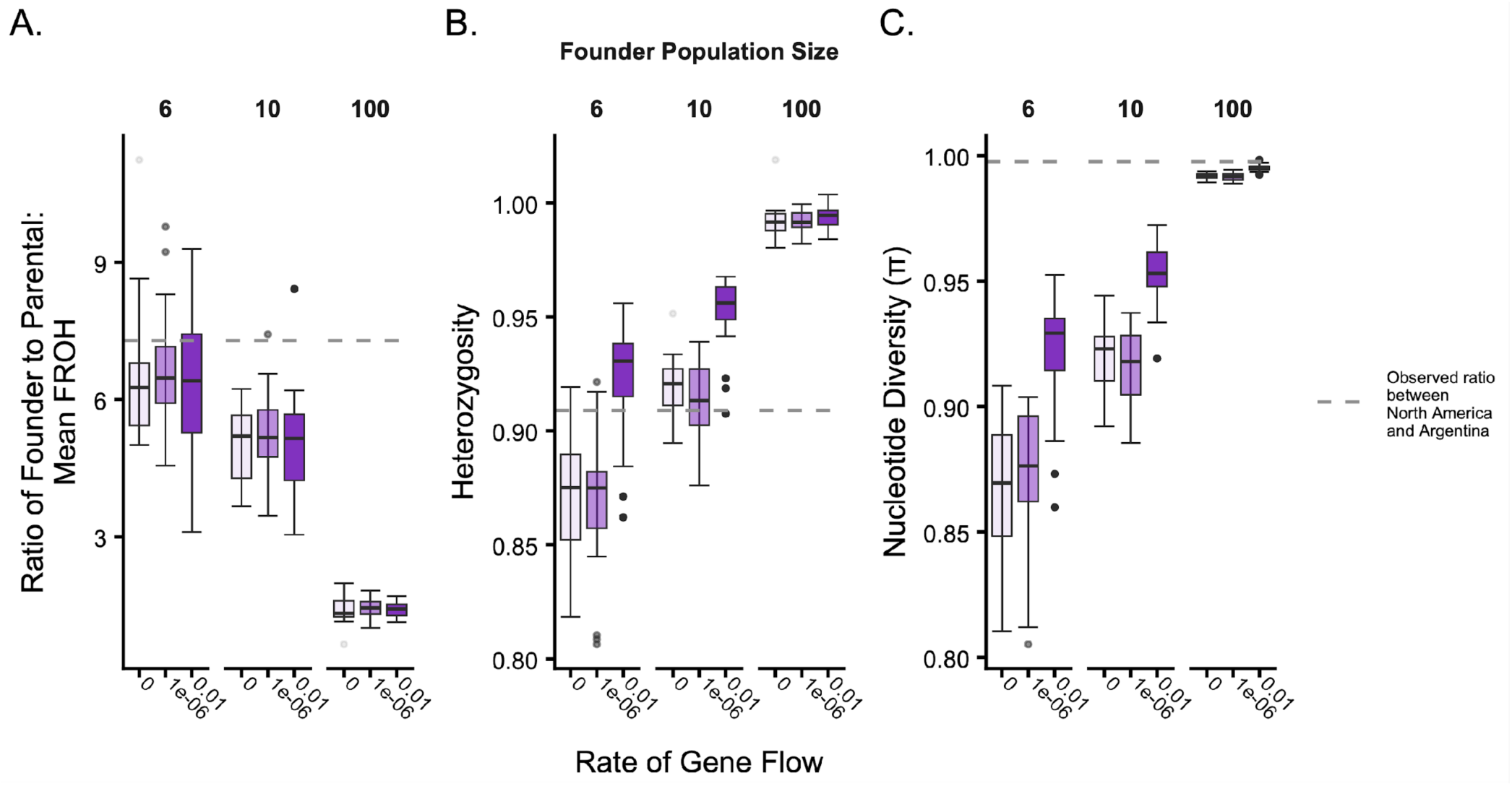
Comparison of population genetic attributes obtained through simulated data with varying conditions of founder population size and gene flow, versus values obtained from the Barn Swallows. **(A)** Ratio of founder to parental mean fraction of the genome in ROH. Darker colors indicate more gene flow. The dotted gray line indicates the observed ratio obtained from North American and Argentine Barn Swallows. **(B)** Ratio of founder to parental heterozygosity. **(C)** Ratio of founder to parental nucleotide diversity.

## Discussion

Switching migratory behavior (i.e., migratory drop offs) appears to promote speciation in birds (*41–43*), and behavioral flexibility without genetic changes has been proposed as a mechanism that promotes migratory switches in birds (*9*, *44*, *45*). We document the initial stages of this process in action happening at ecological timescales, where the process occurs rapidly through the confluence of behavioral, ecological and evolutionary opportunities. In less than five decades, Barn Swallows evolved a new breeding population thousands of kilometers from their original breeding range. We suggest that behavioral flexibility appears to facilitate the initial switch of behavior leading to breeding at the wintering grounds. While slow population growth at the beginning of the colonization episode can provide time for adaptation and accommodation in new environments, Barn Swallows have instead expanded in space and increased in abundance with no evidence of ecological constraints. This ecological opportunity has allowed Argentine Barn Swallows to rapidly expand, maintain genetic diversity, and overcome the likely costs of inbreeding.

Individuals of Argentine Barn Swallows exhibit inbreeding levels comparable to those in populations that have experienced severe declines and inbreeding depression (*46*, *47*), yet they appear to be thriving. At the population level, Argentine Barn Swallows include individuals with high levels of inbreeding as well as admixed individuals, which may buffer the negative effects of inbreeding in combination with rapid population growth. Barn Swallows, as a cosmopolitan species, have undergone multiple colonization and expansion events on different continents during their evolutionary history (*48*). These likely involved broad population size fluctuations, potentially with purging of deleterious variation, which can influence the probability of persistence after founder events (*6*). Indeed, a new population of non-migratory Barn Swallows has recently been described in India, exhibiting the behavioral flexibility of this colonizer species (*49*). Here we have shown that rapid exponential population growth may be a powerful mechanism promoting establishment and persistence following founder events in new breeding localities, just as has been recently found to be determinant for populations recovering from *in situ* bottlenecks (*50*).

## Acknowledgments

We thank Trevor Price, Nicholas Bayly, and Rebecca Safran for valuable feedback during the conception of the project. We are grateful to members of the Toews lab, the Winger Lab, Jake Socha, and Matthias Steinruecken for helpful discussions, and to the members of the Gómez-Bahamón Lab and Trevor Price for comments on earlier drafts of the manuscript. We are also grateful to Pablo Brandolín and Eli Bridge for providing fieldwork support in Cordoba. We thank the curators and collection managers at the American Museum of Natural History, the Field Museum of Natural History, and the Museum of Vertebrate Zoology at the University of California, Berkeley, for their assistance with tissue loans. We thank Anna Maria Calderón, Stephanie Szarmach, Roberto Márquez, Teresa Pegan and Matthew Armstrong for advice on analyses. Portions of this research were conducted with computational resources provided by The Institute for Computational and Data Sciences at The Pennsylvania State University (https://ICDS.psu.edu). We also acknowledge Advanced Research Computing at Virginia Tech for providing computational resources and technical support that have contributed to the results reported within this paper (URL: https://arc.vt.edu/). Finally, we thank Joaquín Cereghetti for showing Valentina a nest of Barn Swallows in a culvert at La Pampa Province of Argentina while studying Fork-tailed flycatchers in 2014.

## Author contributions

Conceptualization: VGB, DPLT, JIA, ZAS

Data collection: VGB, JIA, FG, CT, MP

Analyses: VGB, JS, MP

Funding acquisition: VGB, JIA, DPLT, ZAS

Project administration: VGB, DPLT

Supervision: DPLT, JIA

Writing – original draft: VGB

Writing – review & editing: VGB, DPLT, JIA

## Data and materials availability

All sequence data generated in this study will be available on SRA upon acceptance of the manuscript. Likewise, code and metadata used to perform the analyses presented in this study will be published in Figshare upon acceptance of the manuscript.

## Materials and Methods

### Range expansion and abundance

To estimate the range expansion of the breeding population of Barn Swallows in Argentina we first searched the literature for natural history notes on google scholar that documented breeding populations of Barn Swallows in southern South America. Additionally, we searched Google Scholar we searched using the terms “Barn swallow breeding population in Argentina” and “Barn swallow breeding population in South America” in English and Spanish. We also searched the citizen science database *eBird* (*29*) for comments and observations on active nests of Barn Swallows in southern South America, which provided information on a nesting locality in Chile. We used the reports that included the coordinates where active nests were observed to estimate the smallest convex polygon that encompassed the localities (convex hull) and the smallest polygon that adapted to the distribution of the points (concave hull), connecting the localities where active nests were reported, using the *R* functions *st_convex_hull* and *concaveman* in the *R* packaged *sf* (*52, 53*) and *concaveman* (*54*), respectively. To calculate the total area, we projected the polygons to an equal-area coordinate reference system (EPSG:6933) to avoid distortions associated with geographic coordinates, and then converted the resulting values from square meters to square kilometers by dividing by 1*e^6^*.

To estimate population growth rates in Argentina, we identified a locality where Barn Swallows had not been reported nesting until the year 2018 (*19*). From 2018 onwards we inspected 23 culverts located between the towns of Achiras and Sampacho, Cordoba, for active nests (i.e., those that had eggs or nestlings that were incubated and fed by adults, respectively) (supplementary Fig. S1). Nest counts were performed every two weeks during the entire breeding season for five years (September to February of 2018-2023), and the total count of active nests included double broods. To estimate the rate of population growth at this locality, we fit an exponential curve in *R* finding that the rate of increase was *r* ∼ 0.58 (unit: active nests). To estimate the population growth rate of North American breeding Barn Swallows we used the publicly available and curated data from the North American Breeding Bird Survey (*32*, *33*), and this data includes estimations of relative abundance using hierarchical models (*33*). We estimated the rate of decline since 1980 using the same exponential function described above, finding a rate of a decline of *r* ∼ −0.015.

### Sampling

We captured birds at three localities in Argentina; two in Cordoba Province and an additional site near the center of colonization in Buenos Aires Province (supplementary Table 1). We located breeding colonies by inspection of nests under small bridges and culverts. Once active nests where identified, we placed a mist net at one end of the structure and then one researcher walked inside the structure from the opposite side. We then startled the birds, which flew towards the mist net. We placed individual birds in cotton bags and processed them first by marking them with a metal band with a unique identifying number on one of their tarsi and then measuring them. We then extracted a blood sample from the brachial vein of each individual and sexed them based on plumage characteristics when possible (*55*), as well as on evidence of brood patch in females and cloacal protuberance in males. We released all birds after these procedures.

To obtain samples from North America, we requested museum tissue loans from birds collected during the breeding season only (June and July) from the American Museum of Natural History (Eastern North American tissues, supplementary Table 1), the Field Museum of Natural history (Tissues in the Midwest, supplementary Table 1), and the Museum of Vertebrate Zoology, Berkeley (Western North American tissues, supplementary Table 1). We additionally included sequences from a published dataset from the Rocky Mountains in Colorado, but not for ROH analysis because depth coverage in these sequences was <10X (*34*)(SRA: SRS33028, supplementary Table. 1).

### DNA extraction and Library preparation

We extracted DNA using the Qiagen DNeasy Blood and Tissue Kit and protocol. For samples collected in the field, we extracted DNA from 75μL of blood. For museum tissues, we extracted DNA from 10-25mg of tissue. We then prepared libraries using an Illumina DNA Prep (M) Tagmentation (96 samples) kit and protocol. Next, we indexed the samples with Nextera DNA CD indexes, and pooled libraries and sequenced them with Illumina NextSeq500 at the Huck Institutes of Life Sciences Genomics Core, Penn State University. We completed whole genome resequencing for 30 individuals resulting in a mean depth coverage of ∼6X, supplementary Table 1) and re-sequenced 9 of those individuals that had lower than ∼11X depth coverage and included individuals that had >10X depth coverage from the first run and used these data for ROH and outlier region analyses (supplementary Table 1).

### Filtering and Genotyping

We aligned whole-genome resequencing reads to a published chromosome-level annotated reference genome of the Barn Swallows (*Hirundo rustica*; PRJNA636192; Vertebrate Genomes Project). To do this, we first indexed the genome in *Bowtie2* (*56*) and then we aligned the whole genome resequencing reads to it (also using *Bowtie2*), with the *very-sentive-local* preset and assuming *Phred+33* quality score encoding. We converted SAM files to BAM files and sorted them in *SAMtools 8.3.1* (*57*). We marked and masked PCR duplicates using *Picard Tools 2.20.8* (*58*), and the resulting BAM files were indexed using *SAMtools 8.3.1*. We inspected read quality and statistics using *Qualimap2.3* (*59*).

To study population differentiation, we estimated genotype likelihoods in *ANGSD* (*60*) generating a SAF file for western and eastern North America, and Argentina, with log-likelihood ratios of allele frequencies for each site. We then generated a folded site frequency spectrum and weighted *F*_ST_ using *realSFS* also in *ANGSD.* We performed a windowed analysis of 10kb non-overlapping windows. To compare observed *F*_ST_ to a null expectation, we generated three randomized groups sampled from pairwise combinations of the Argentine, Eastern North American, and Western North American samples. We then repeated our sliding window *F*_ST_ calculation across the genome between each of these randomized groups and the group that was excluded. We iterated this process 500 times to generate our null distribution. We plotted this distribution along the genome, displaying the interval containing 95% of the null distribution. We then excluded the *Z* chromosome from population differentiation analyses to avoid biases due to differences in sex sampling between males and females. We ran a principal components analysis in *ANGSD*, followed by an admixture analysis in *NGSadmix* (*61*) for K=2-4, and a genetic distance analysis in *ngsDist* (*62*). A topology using the genetic distance analyses was generated using *FastME 2.0* (*63*).

### Heterozygosity and genetic diversity

We estimated the site frequency spectrum using *ANGSD* and from this obtained genetic diversity (*Watterson’s θ*), and nucleotide diversity (*θπ*). Specifically, we estimated *θπ* and *Watterson’s θ* with *realSFS*, and we estimated Tajima’s D using *thetaStat*. To facilitate comparisons among populations, we divided all genetic diversity metrics by the number of sites. We then estimated individual heterozygosity also using *ANGSD* with genotype likelihoods on a per-individual basis. The site frequency spectrum was then inferred using *realSFS*, which we subsequently used for visualization and downstream analyses in *R.* We excluded the sex chromosome *Z* from these analyses as well.

### Calling runs of homozygosity

To call runs of homozygosity (ROH) we analyzed the data with mean depth coverage ∼11X, resulting in data for 8 individuals (supplementary Table 1). First, we performed variant calling across all individuals in *GATK3* (*64*) generating files in a genomic variant call format. The resulting files were indexed in *BCFtools* (*65*) and site quality was filtered with *Phred-scaled* quality score of 50 or greater. We did all the downstream analyses described below using *BCFtools*, unless otherwise stated. We filtered out sites with combined read depth across individuals that were less than 80 or greater than 193 (these values correspond to 5th and 95th percentiles for total depth of our samples, respectively). We excluded multi-nucleotide polymorphisms, indels, complex variants, sites with more than one alternate allele, and sites with more than two missing genotypes. We then generated a subset vcf file per population with *GATK*3 and indexed them with *BCFtools.* We excluded sites with excess heterozygosity per population (i.e., defied as sites with 75% or more heterozygote genotypes). We calculated Depth (DP) thresholds using a *python* script. We inspected the distribution of DP thresholds in *R* and filtered individual phenotypes. Next, we filtered for genotype quality based on GQ scores. Then we generated a dummy centromere file with 0s and 1s and filtered sites by population based on missing GT thresholds. If the site had more than 3 missing genotypes, we filtered it out. After indexing, we filtered out the *Z* chromosome and removed scaffolds. We excluded one individual (B15500) due to low data after filtering, resulting in 7 remaining samples. We created a *tgls* file using a *python* script and used *PLINK 1.9* (*66, 67*) to create the *tped* and *tfam* files. Finally, we called ROH in *Garlic* (*68*) using a window size of 40. We discarded ROH smaller than 0.5mb.

To estimate the fraction of the genome in ROH, we estimated genome mappability using *genmap* (*69*), with a *kmer* size of 100 and error rate of 2 (Fig. S5). We then filtered for regions of the genome with a mappability score of 1 and summed the length of these regions to get the mappable genome size (0.909 Gb). We divided the lengths of the summed ROH of each size category by the estimated mapsize to get FROH. Inbreeding coefficients were estimated as the correlation between uniting gametes (Fhat3) using *PLINK* with the option --ibc on autosomal SNPs pruned for high linkage disequilibrium.

### Fastsimcoal

Demographic history was inferred using the composite-likelihood approach implemented in *fastsimcoal2* (*70*). We first generated a two-dimensional joint site frequency spectrum in *ANGSD* and reformatted in *R* as a matrix of allele frequency counts for each population. We evaluated six alternative demographic models, including scenarios with bidirectional migration, unidirectional migration, and no migration, with and without population growth. For each model, we performed parameter estimation by fitting simulated SFS to the observed 2D-SFS using a maximum likelihood framework. Each likelihood estimation was based on 100,000 coalescent simulations (-n 100000) and 150 conditional maximization cycles (-L 150). We estimated model parameters, including effective population sizes, divergence time, migration rates, and population growth. We run each demographic model multiple independent times to ensure convergence, with 80 replicates per model for migration-only scenarios and 20 replicates per model for models including population growth. Model support was evaluated using Akaike Information Criterion (AIC), estimated from maximum likelihood estimates and number of parameters in each model. Rather than selecting a single best-supported model, we visualized AIC values across runs in *R* and highlighted the highest-ranking models within parameter space to assess support across alternative demographic scenarios.

### F_ST_ outliers, Tajima’s D, and Nucleotide Diversity

We analyzed allelic frequency differentiation from North and South American breeding swallows by using the ∼11X coverage data and genotype likelihoods in *ANGSD*. We performed all analyses below in *ANGSD,* unless otherwise stated. We estimated the joint site frequency spectra between populations using *realSFS* with iterative optimization (-maxIter 300). For analyses requiring folded spectra, we folded the SFS prior to downstream inference. We then estimated genome-wide genetic differentiation (F*_ST_*) between populations using *realSFS fst*, followed by window-based estimation using *realSFS fst stats2*. Finally, we estimated per window F*_ST_* in non-overlapping sliding windows of 10 Kb.

To characterize genome-wide patterns of diversity and neutrality, we estimated per-site θπ using *realSFS*, as well as π and Tajima’s D. We summarized these statistics in non-overlapping 10 kb windows. We plotted genome-wide patterns of differentiation using the *manhattan* function in the *qqman* package (*71*) in *R*. Regions of elevated genetic differentiation were identified from F*_ST_* scans and further examined in the genome browser *Geneious Prime 2025.1* to determine their genomic context and proximity to annotated genes. These regions were then compared with patterns of nucleotide diversity and Tajima’s D to assess signatures of selection.

### Simulations

We performed simulations using *msprime* (*72*), a coalescent-based simulator that models genetic variation using tree sequence recording (i.e., the ancestral genealogic history). We estimated summary statistics using *tskit* (*73*, *74*). We simulated the demographic history of North American and Argentine Barn Swallow populations based on parameter estimates obtained from *fastsimcoal2* and the population growth rates estimated with empirical and citizen science data. The North American population was initialized with an effective population size of 3.2*e^4^* and a growth rate of −0.015, whereas the Argentinian population was initialized with an effective population size of 3.1e8 and a growth rate of 0.58. Growth rates follow the backward-time parameterization implemented in *msprime*. We set a population split occurring 30 generations in the past. We simulated 20 Mb genomic regions with a recombination rate of 3.1 × 10⁻⁸ and a mutation rate of 4.42 × 10⁻⁹ per site per generation based following estimates from Collared flycatchers (*39*, *75*). For each simulation, we sampled 40 diploid individuals per population. Simulations were repeated 20 times for each parameter combination.

From the resulting simulated tree sequences, we calculated individual heterozygosity and nucleotide diversity *π* using site-based diversity statistics. We estimated individual heterozygosity by averaging per-individual diversity across all sampled individuals within each population. We generated variant call format files from simulated data for downstream analyses, including estimation of runs of homozygosity (ROH) following the same pipeline described above. We estimated FROH across ROH length classes. Summary statistics were then aggregated across parameter combinations to evaluate the effects of gene flow and ancestral population size on ROH patterns. For graphic purposes we converted all metrics to ratios of founder to parental populations, to make them comparable to the ratio of the South American to North American populations that we compared them to.

