## Supplementary_figures for "Rapid expansion overcomes inbreeding in cross-hemisphere colonization of Barn Swallows"

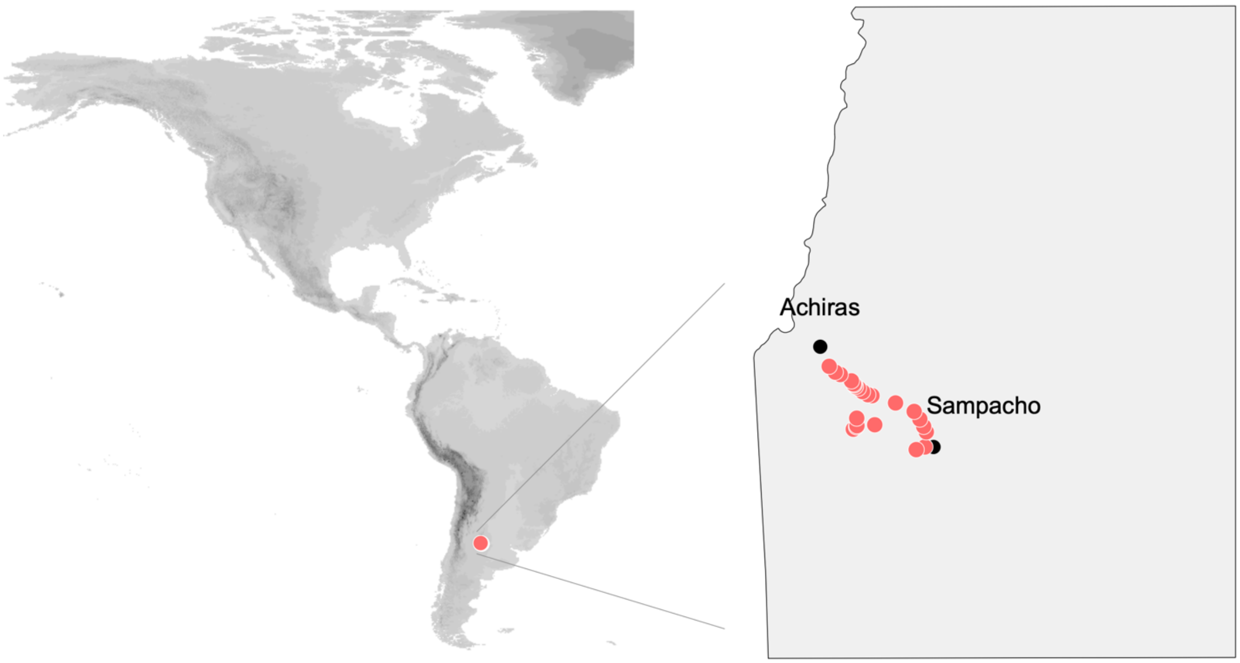


**Fig. S1. Sites where active Barn Swallow nests were monitored.** The red dots indicate the 23 bridges and culverts that were monitored during five consecutive years between the towns of Sampacho and Achiras. The black dots indicate town centers.

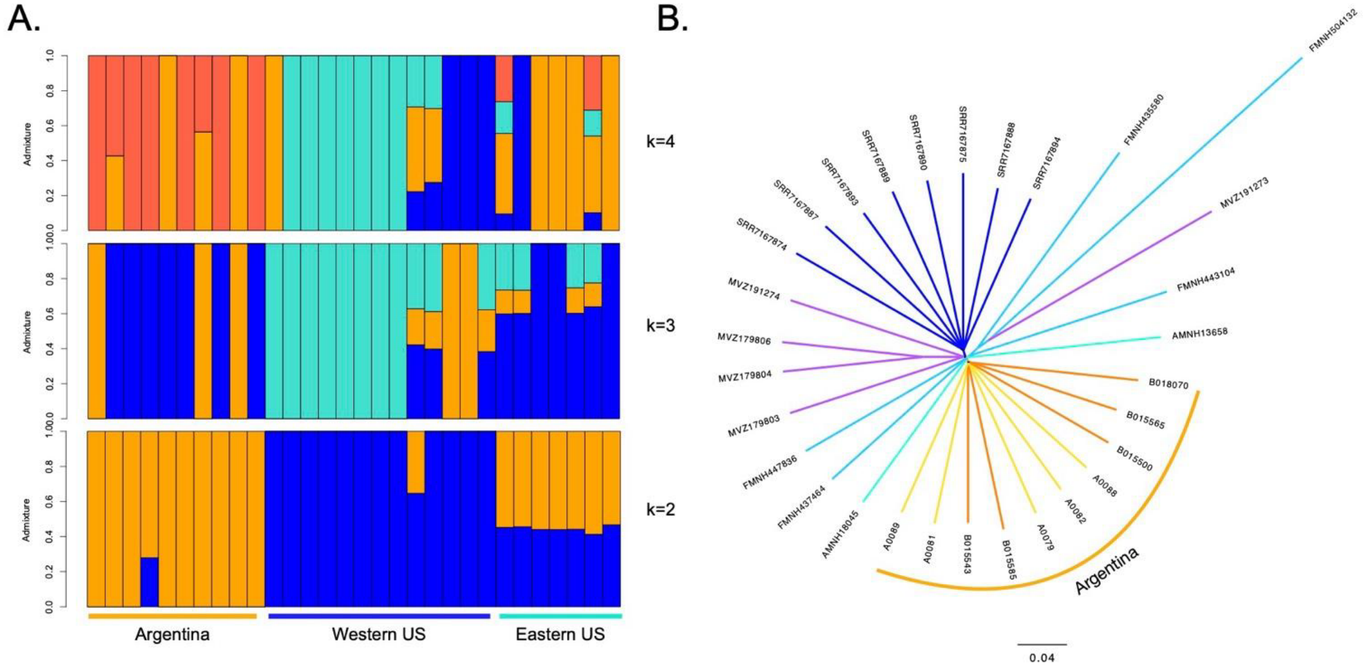


**Fig. S2. Population structure of Barn Swallows. A)** Admixture proportions of individuals and cluster assignment when considering 2, 3 and 4 groups. **B)** Distance tree showing that Argentine Barn Swallows form a monophyletic group. Dark blue indicate samples from Colorado, purple from western north America and aquamarines from eastern North America.

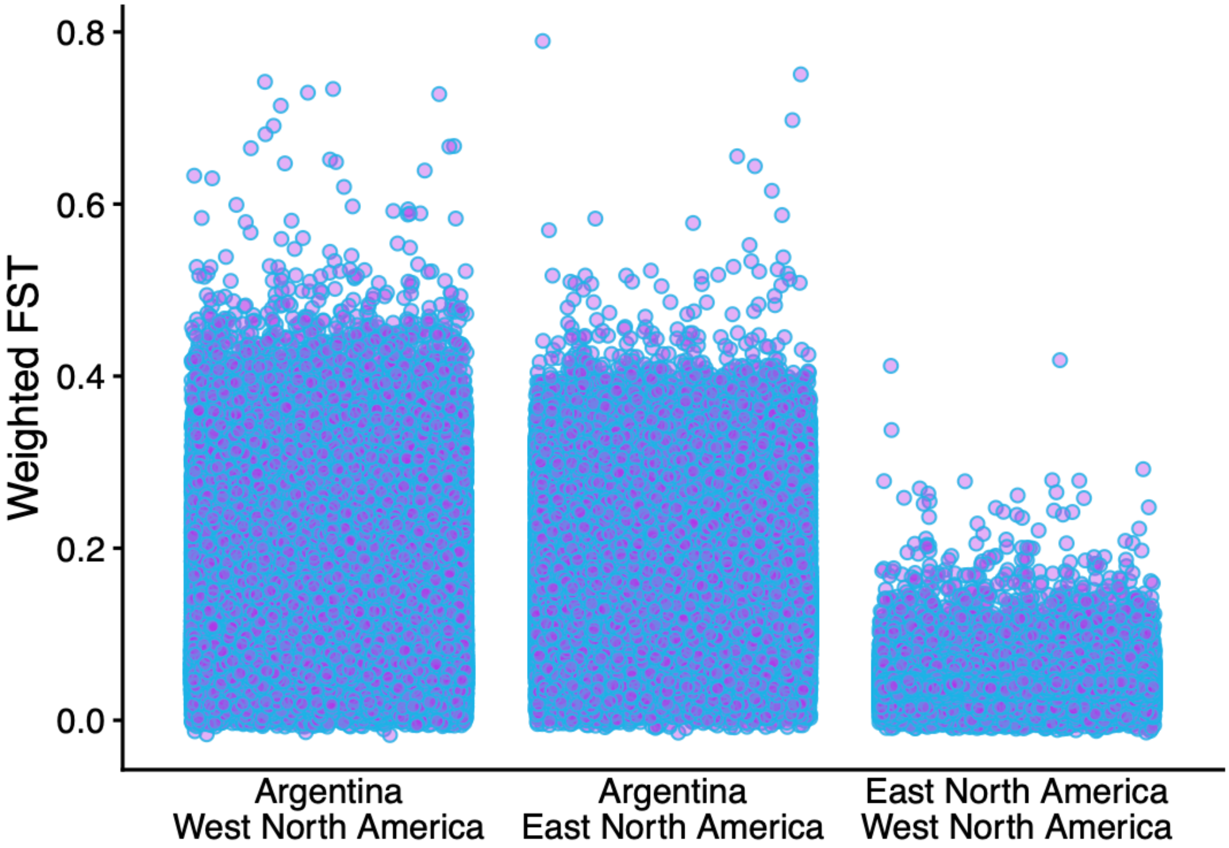


**Fig. S3. Genetic differentiation across the genome in Barn Swallows.** Genetic differentiation estimated with weighted $F_{ST}$ across 10 Kb non-overlapping windows for pairwise comparisons between eastern North American breeding birds (including from the Midwest region of the United States of America) with South American breeding birds, western North American breeding birds with South American breeding birds, and between eastern and western North American breeding birds.

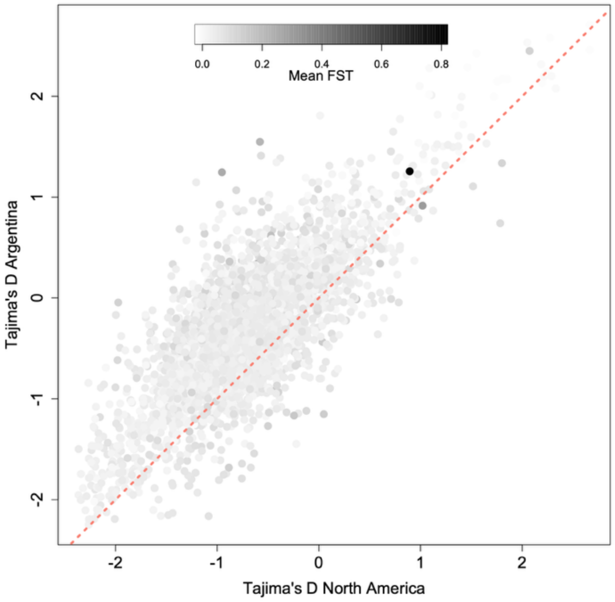


**Fig. S4. Evidence of balancing selection on chromosome 6 of Barn Swallows.** Tajima's D estimates for North America and Argentina across chromosome 6, where positive values indicate evidence of balancing selection and negative values of selective sweeps. Outlier regions are colored in black corresponding to the estimated $F_{ST}$ values.

The points closer to the red dotted line indicate evidence for similar processes action on that region in Argentina and North America.

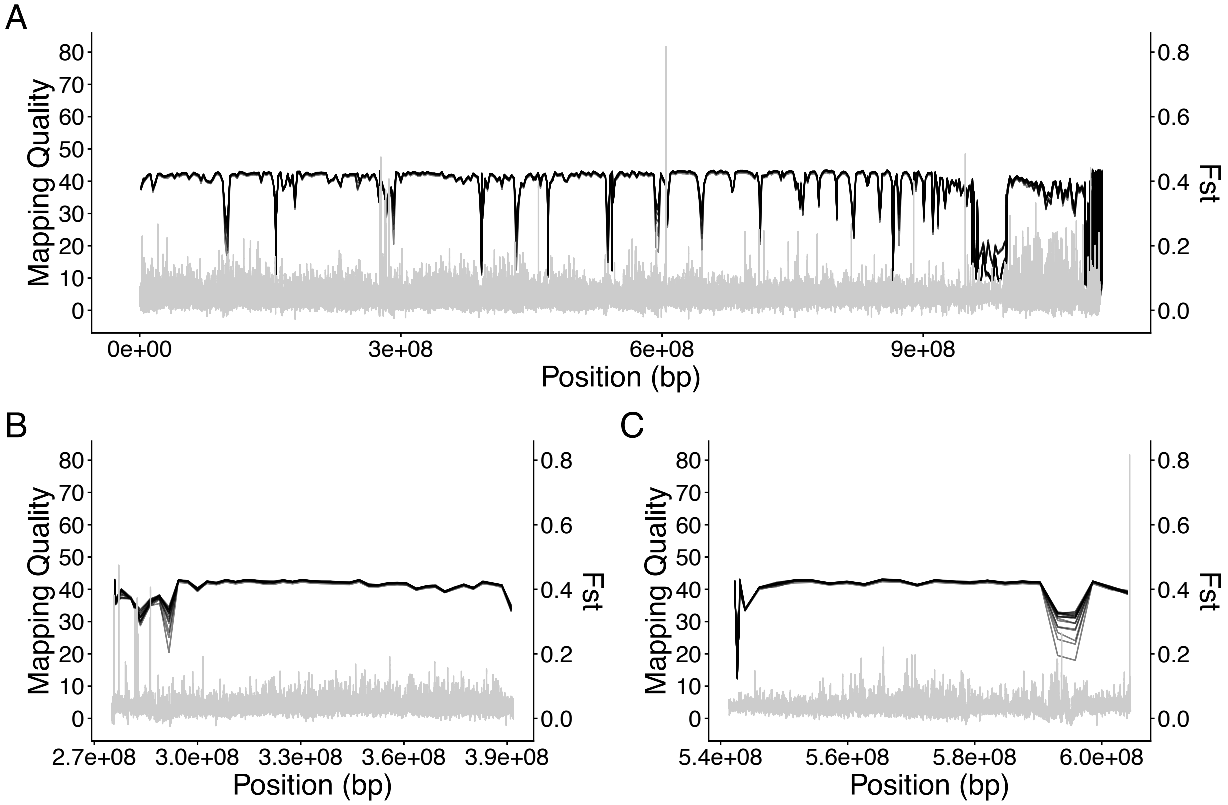


**Fig. S5. Mappability across the Barn Swallow genome and the two chromosomes containing regions of elevated genetic differentiation.** (A) Whole-genome mappability in black with $F_{ST}$ in gray. (B) Mappability across chromosome 3 in black with $F_{ST}$ in gray. (C) Mappability across chromosome 6 in black with $F_{ST}$ in gray.

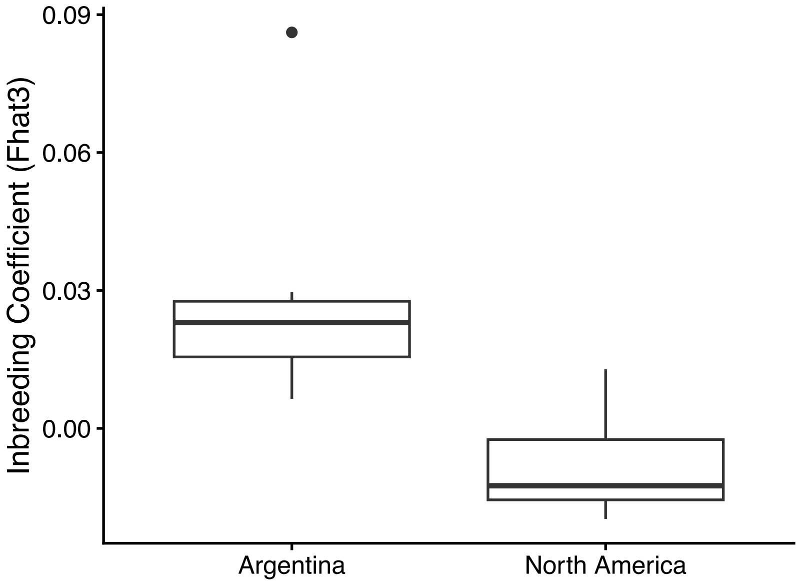


**Fig. S6.** Individual inbreeding coefficients (Fhat3) for Argentine and North American breeding Barn Swallows.

**Table S1.** Sampling information and whole-genome sequencing statistics for all Barn Swallow individuals included in this study.

| Sample ID | Year | Source | Sex | Latitude | Longitude | Mapped reads | Percent | Coverage (mean) | SD |
|---|---|---|---|---|---|---|---|---|---|
| B015500 | 2020 | Wild | male | -38.2794 | -58.5803 | 45,413,869 | 97.91% | 6.9819 | 17.7166 |
| B018070 | 2019 | Wild | male | -38.2822 | -58.5814 | 34,830,815 | 98.01% | 5.4637 | 15.5745 |
| B015543 | 2021 | Wild | male | -38.2822 | -58.5814 | 47,978,143 | 97.93% | 7.4873 | 20.8191 |
| A0079 | 2019 | Wild | male | -33.324486 | -64.752667 | 42,511,915 | 97.79% | 6.5422 | 20.7493 |
| A0081 | 2019 | Wild | male | -33.324486 | -64.752667 | 43,371,839 | 97.82% | 6.8069 | 19.5191 |
| A0082 | 2019 | Wild | male | -33.324486 | -64.752667 | 52,581,206 | 97.77% | 7.8161 | 25.1635 |
| A0089 | 2019 | Wild | male | -33.292167 | -64.810658 | 40,435,136 | 97.87% | 6.2532 | 19.2833 |
| A0088 | 2019 | Wild | male | -33.292167 | -64.810658 | 46,366,447 | 97.79% | 7.1109 | 24.4631 |
| B015565 | 2020 | Wild | male | -38.2794 | -58.5803 | 67,346,823 | 97.94% | 10.0324 | 23.1615 |
| B015585 | 2020 | Wild | male | -38.2794 | -58.5803 | 44,783,265 | 98.24% | 6.8542 | 18.9843 |
| SRR7167875 | NA | NCBI | female | 40.0274 | -105.2519 | 38,088,726 | 98.20% | 5.9907 | 60.5452 |
| SRR7167874 | NA | NCBI | male | 40.0274 | -105.2519 | 32,534,061 | 98.04% | 5.0448 | 49.0341 |
| SRR7167890 | NA | NCBI | male | 40.0274 | -105.2519 | 40,785,284 | 98.05% | 6.4892 | 70.9304 |
| SRR7167889 | NA | NCBI | female | 40.0274 | -105.2519 | 45,375,154 | 98.18% | 7.2369 | 81.1971 |
| SRR7167888 | NA | NCBI | female | 40.0274 | -105.2519 | 44,184,228 | 98.25% | 7.1509 | 82.1276 |

| | | | | | | | | | |
|---|---|---|---|---|---|---|---|---|---|
| SRR7167887 | NA | NCBI | male | 40.0274 | -105.2519 | 39,078,468 | 98% | 6.0816 | 78.9703 |
| SRR7167894 | NA | NCBI | female | 40.0274 | -105.2519 | 46,428,971 | 98.25% | 7.464 | 73.8174 |
| SRR7167893 | NA | NCBI | female | 40.0274 | -105.2519 | 46,508,316 | 98.20% | 7.379 | 72.8328 |
| A0079 | 2019 | Wild | male | -33.324486 | -64.752667 | 29,538,314 | 97.82% | 4.6586 | 15.368 |
| A0081 | 2019 | Wild | male | -33.324486 | -64.752667 | 36,673,628 | 97.75% | 5.591 | 17.5694 |
| A0082 | 2019 | Wild | male | -33.324486 | -64.752667 | 31,573,443 | 97.75% | 4.8089 | 16.1439 |
| A0088 | 2019 | Wild | male | -33.292167 | -64.810658 | 23,909,194 | 97.70% | 3.3825 | 12.3179 |
| A0089 | 2019 | Wild | male | -33.292167 | -64.810658 | 28,038,595 | 97.92% | 4.2514 | 12.5402 |
| AMNH13059 | 2002 | AMNH | NA | 40.770278 | -73.153611 | 26,529,081 | 97.83% | 3.9743 | 38.066 |
| AMNH13656 | 2004 | AMNH | NA | 40.770278 | -73.153611 | 28,067,311 | 97.65% | 4.434 | 52.6756 |
| AMNH13657 | 2004 | AMNH | NA | 40.901389 | -73.343056 | 31,710,003 | 97.80% | 5.0045 | 61.4587 |
| AMNH13658 | 2004 | AMNH | NA | 40.901389 | -73.343056 | 25,723,452 | 97.36% | 3.8013 | 38.2579 |
| AMNH18045 | 2008 | AMNH | NA | 41.451389 | -71.524167 | 33,081,442 | 96.87% | 5.2898 | 37.614 |
| B015500 | 2020 | Wild | male | -38.2794 | -58.5803 | 33,345,570 | 97.77% | 4.5011 | 12.4071 |
| B015543 | 2021 | Wild | male | -38.2822 | -58.5814 | 27,347,234 | 97.92% | 4.3022 | 12.7804 |
| B018070 | 2019 | Wild | male | -38.2822 | -58.5814 | 30,899,806 | 97.89% | 4.6902 | 14.2611 |
| FMNH435580 | 2002 | FMNH | NA | 41.750278 | -87.919444 | 23,063,145 | 97.64% | 3.4209 | 19.9235 |
| FMNH437464 | 2000 | FMNH | NA | 43.899444 | -89.5 | 30,061,209 | 96.41% | 4.7638 | 23.7654 |
| FMNH443104 | 2003 | FMNH | NA | 41.422778 | -87.986111 | 23,560,737 | 96.41% | 3.661 | 13.2603 |
| FMNH477836 | 2010 | FMNH | NA | 41.871389 | -87.634167 | 29,300,228 | 96.16% | 4.5773 | 16.3728 |
| MVZ179803 | 1996 | MVZ | NA | 37.909167 | -122.686111 | 29,832,286 | 96.27% | 4.6208 | 26.2258 |
| MVZ179804 | 1996 | MVZ | NA | 37.933333 | -122.737778 | 31,997,848 | 97.22% | 4.9901 | 25.9767 |
| MVZ179806 | 1996 | MVZ | NA | 37.933333 | -122.737778 | 30,736,649 | 96.77% | 4.738 | 21.5481 |
| MVZ191274 | 2007 | MVZ | NA | 41.225278 | -121.1425 | 30,452,619 | 96.85% | 4.7478 | 22.241 |
| A0079 | 2019 | Wild | male | -33.324486 | -64.752667 | 72,050,172 | 97.80% | 11.2008 | 35.8836 |
| A0081 | 2019 | Wild | male | -33.324486 | -64.752667 | 80,045,508 | 97.79% | 12.3979 | 36.7628 |
| A0082 | 2019 | Wild | male | -33.324486 | -64.752667 | 84,154,739 | 97.76% | 12.625 | 41.0481 |
| A0089 | 2019 | Wild | male | -33.292167 | -64.810658 | 68,473,735 | 97.89% | 10.5047 | 31.5374 |
| A0088 | 2019 | Wild | male | -33.292167 | -64.810658 | 70,275,637 | 97.76% | 10.4934 | 36.5381 |
| AMNH13658 | 2004 | AMNH | NA | 40.901389 | -73.343056 | 95,961,567 | 97.21% | 14.9806 | 145.6879 |

| | | | | | | | | | |
|---|---|---|---|---|---|---|---|---|---|
| AMNH18045 | 2008 | AMNH | NA | 41.451389 | -71.524167 | 100,676,590 | 96.47% | 15.9662 | 114.9356 |
| B015500 | 2020 | Wild | male | -38.2794 | -58.5803 | 33,324,616 | 97.77% | 4.4983 | 12.3999 |
| B015543 | 2021 | Wild | male | -38.2822 | -58.5814 | 75,325,292 | 97.93% | 11.7895 | 33.3278 |
| B018070 | 2019 | Wild | male | -38.2822 | -58.5814 | 65,730,593 | 97.96% | 10.1539 | 29.5492 |
| B015565 | 2020 | Wild | male | -38.2794 | -58.5803 | 67,346,823 | 97.94% | 10.0324 | 23.1615 |
| FMNH435580 | 2002 | FMNH | NA | 41.750278 | -87.919444 | 108,728,250 | 97.28% | 16.4946 | 104.444 |
| FMNH437464 | 2000 | FMNH | NA | 43.899444 | -89.5 | 110,602,550 | 95.86% | 17.2708 | 82.8867 |
| FMNH443104 | 2003 | FMNH | NA | 41.422778 | -87.986111 | 108,311,345 | 95.62% | 16.4996 | 64.7031 |
| FMNH504132 | 2015 | FMNH | NA | 44.899 | -89.567 | 81,299,387 | 95.11% | 12.6015 | 54.3249 |
| MVZ179804 | 1996 | MVZ | NA | 37.933333 | -122.737778 | 105,451,974 | 96.99% | 16.3301 | 88.0606 |
| MVZ179806 | 1996 | MVZ | NA | 37.933333 | -122.737778 | 102,415,256 | 96.69% | 15.9943 | 74.3069 |
| MVZ191273 | 2007 | MVZ | NA | 40.934 | -120.205 | 90,145,911 | 97.02% | 14.1 | 69.1777 |
